# The origin and evolution of the gonococcal beta-lactamase plasmid, and implications for public health

**DOI:** 10.1101/2025.01.03.631176

**Authors:** T. A. Elsener, A. Cehovin, C. Philp, K. Fortney, S.M. Spinola, M.C.J. Maiden, C.M. Tang

## Abstract

*Neisseria gonorrhoeae* is a leading cause of sexually transmitted infection (STI) and a priority AMR pathogen. Two narrow host range plasmids, p*bla* and pConj, have contributed to ending penicillin and tetracycline therapy, respectively, and undermine current prevention strategies including Doxy-PEP. Here, we investigated the origin and evolution of the beta-lactamase plasmid, p*bla*. We show that the interplay between p*bla* and pConj influences their co-occurrence and the spread of p*bla* in the gonococcal population. We demonstrate that p*bla* was acquired by the gonococcus on at least two occasions from *Haemophilus ducreyi*, and describe the subsequent evolutionary pathways taken by the three major p*bla* variants. Changes that mitigate fitness costs of p*bla* and the emergence of TEM beta-lactamases which confer increased resistance have contributed to the success of p*bla*. In particular, TEM-135, which has arisen in certain p*bla* variants, increases resistance to beta-lactams and only requires one amino acid change to become an extended spectrum beta-lactamase (ESBL). The evolution of p*bla* underscores the threat of plasmid-mediated resistance to current therapeutic and preventive strategies against gonococcal infection. Given the close relationship between p*bla* and pConj, widespread use of Doxy-PEP is likely to promote spread of pConj and p*bla*, and emergence of plasmid-mediated ESBL in the gonococcus, with dire public health consequences.

## INTRODUCTION

*Neisseria gonorrhoeae* causes ∼80 million sexually transmitted infections (STIs) annually^4^ and is a WHO priority pathogen due to its extensive antimicrobial resistance (AMR)^5^. *N. gonorrhoeae* has two resistance plasmids, pConj and p*bla*. These plasmids contributed to the cessation of tetracycline and penicillin therapy, and undermine doxycycline post-exposure prophylaxis (Doxy-PEP)^6,7^. pConj and p*bla* are highly prevalent in low and middle-income countries (LMICs) where syndromic treatment of STIs with doxycycline has been recommended^1,7,8^. Therefore, it is important to understand the factors driving the success of these plasmids in gonococcal populations.

pConj is a 39-42 kb conjugative plasmid, that can mediate tetracycline resistance^7,10^, and can be categorised into seven variants^8^. p*bla* emerged in the gonococcus in the 1970s and encodes the TEM beta-lactamase conferring penicillin resistance^11,12^. p*bla* TEM beta-lactamases require one or two amino acid changes to become an extended-spectrum beta-lactamase (ESBL)^13^, which would render third-generation cephalosporins, the current first-line treatment, ineffective^14^. p*bla* usually is closely associated with pConj which can mobilise p*bla*^8,15^.

There are three main p*bla* variants, characterised by distinct gene presence/absence patterns^1^. The 7.4 kb p*bla*.2 (p*bla* Asia) has been considered the ancestral plasmid^16^. p*bla*.1 (5.6 kb, p*bla* Africa) has a deletion in the replication region, while p*bla*.3 (5.1 kb, p*bla* Rio) lacks the region implicated in p*bla* mobilisation^1,16^. Variants of p*bla* are associated with certain pConj variants and TEM alleles^1^. p*bla*.1 mostly carries TEM-1 or TEM-1_P14S_, while p*bla*.3 is associated with TEM-135; p*bla*.2 carries TEM-1 or TEM-135. Importantly, the M182T substitution in TEM-135 is a ‘stepping stone’ mutation before becoming an ESBL^13,17^.

Here, we investigated evolution and characteristics of the p*bla* variants. We demonstrate that the spread and distribution of p*bla* in gonococci results from the dynamic interplay of its intimate association with pConj, fitness costs, and resistance levels. p*bla* has evolved to avoid fitness costs and confer higher resistance to beta-lactams. Our results underline the threats posed by p*bla* and pConj, particularly through the widespread implementation of Doxy-PEP.

## RESULTS

### p*bla* has been acquired at least twice by the gonococcus from *Haemophilus*

*Haemophilu*s spp. harbours plasmids related to p*bla*^18,19^. Therefore, we interrogated *Haemophilus* whole genome sequences (WGS, 4,620 isolates, 12 species, Supplementary Table 1) for p*bla*. We searched for Tn*2* as p*bla* TEM-1b is located on this transposon^20,21^, and confirmed the presence of p*bla* replicons by searching for NEIS2960, NEIS2358, and NEIS2961^1^. Tn*2* is present in 10.2% of *Haemophilus influenzae* (4,337 isolates), with a p*bla*-like plasmid only co-occurring in one *H. influenzae* (PubMLST id: 23482).

In contrast, 22.6% of *H. ducreyi* (7/31 isolates) harbour TEM-1 containing p*bla*-like plasmids; resequencing of *H. ducreyi* HD183 and DMC64^22^ identified distinct 9.1 and 10.9 kb plasmids, respectively. We aligned p*bla*.1 and p*bla*.2 to the *H. ducreyi* plasmids^22^. p*bla*.1 and p*bla*.2 are highly related to the 9.1 and 10.9 kb *H. ducreyi* plasmids, respectively (Figure 1A). Tn*2* is intact in *H. ducreyi* while gonococcal p*bla* lack *tnpA* and have a truncated *tnpR,* with distinct *tnpR* alleles on p*bla*.1 is and p*bla*.2/3, indicating that p*bla*.1 and p*bla*.2 were acquired independently by the gonococcus.

**Figure 1:**
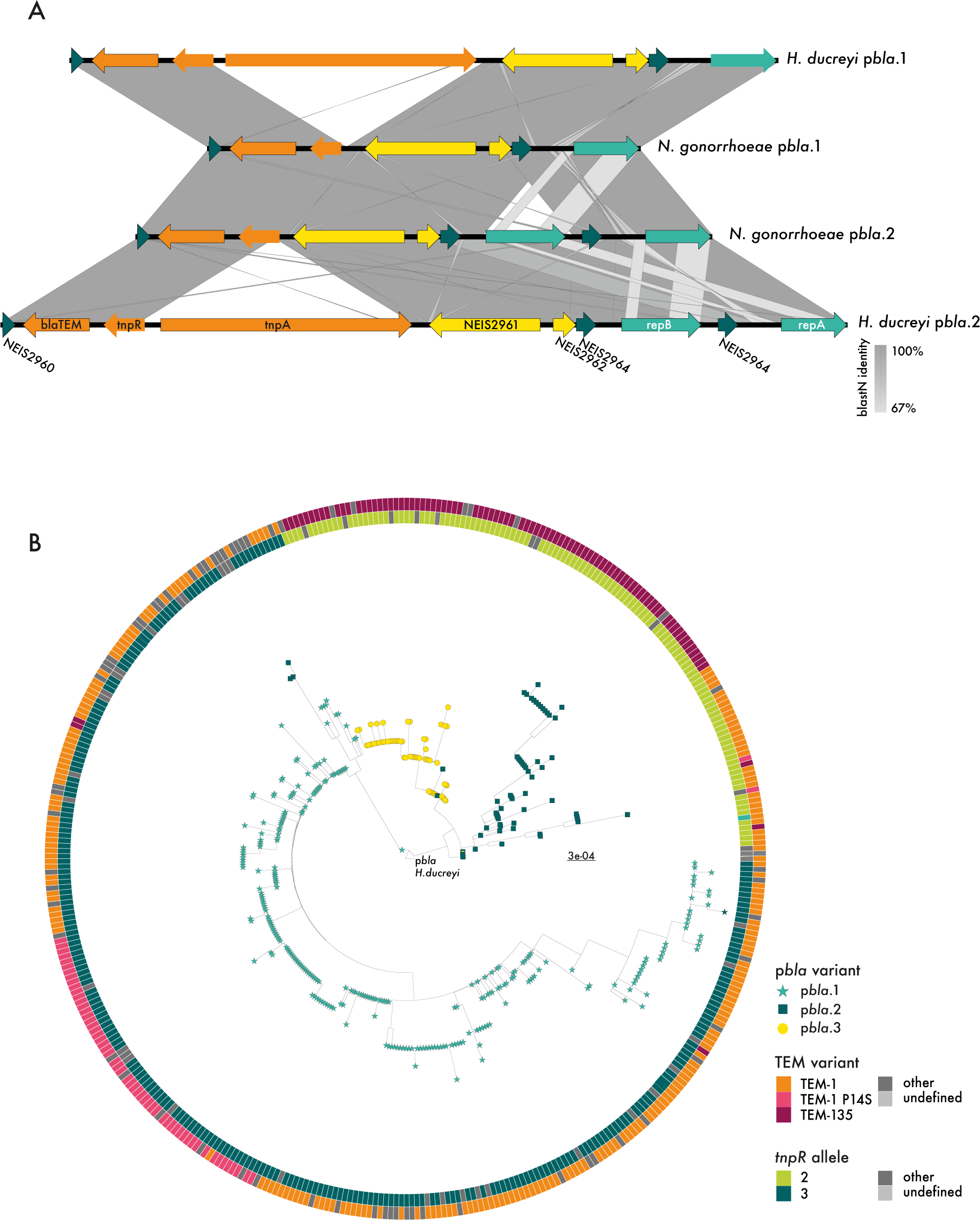
Origin and evolution of p*bla.* (A) Pairwise alignments of p*bla* variants in *H*. *ducreyi* and *N. gonorrhoeae* generated with Easyfig^2^. p*bla* ORFs are coloured according to gene products; yellow, mobilisation proteins; orange, Tn*2*-derived genes including *bla*TEM; turquoise, replication initiation proteins; dark green, undefined/other gene function. Sequence identity between loci is depicted in shades of grey, as indicated. (B) Maximum likelihood tree of 414 gonococcal p*bla* sequences with tips coloured according to p*bla* variant. Circles indicate the *tnpR* allele and the TEM variant carried.

To gain further insights into the evolutionary relationships between p*bla* variants, we examined a subset of p*bla* (414 of 2,758, Supplementary Table 2)^1^ with the 10.9 kb *H. ducreyi* p*bla* as reference; plasmids were from 1979-2022 with the same proportion of variants as the whole population (*i.e.* 70% p*bla*.1, 14% p*bla*.2, 16% p*bla*.3)^1^. Maximum likelihood phylogeny distinguished p*bla* variants into distinct clades, with p*bla*.1 split from the other variants, and p*bla*.3 arising from p*bla*.2 (Figure 1B).

Taken together, gonococcal p*bla*.1 and p*bla*.2 arose by independent acquisition of plasmids expressing TEM-1 from *H. ducreyi*. These events were associated with distinct truncations of Tn*2*, with subsequent emergence of p*bla*.3 from p*bla*.2.

### p*bla* is associated with pConj variants that promote its spread

To understand the association between p*bla* and pConj, we examined the transfer of p*bla* by different pConj variants. Matings were performed between isogenic strains (FA1090 or 2086_K) with Δ*pilD* donors and recipients to prevent transfer by transformation^23-25^. p*bla*.1 was mobilised at a frequency of ∼1% transconjugants/recipient by pConj.1 in both strains (Supplementary Figure 1).

p*bla* is commonly associated with pConj variants 1, 3, 4 and 5 (Figure 2A). Therefore, we evaluated p*bla* mobilisation by different pConj variants. The conjugation frequencies of pConj.1, 3 and 4 were >79% (Figure 2B) but several orders of magnitude lower for pConj.2 and pConj.7 (which are not associated with p*bla*, Figure 2A). The rate of p*bla* mobilisation mirrored conjugation frequencies, with p*bla* transfer by pConj.2 undetectable (Figure 2B), indicating that p*bla* is associated with pConj variants that mobilise it efficiently, and promote its spread in the gonococcal population.

**Figure 2:**
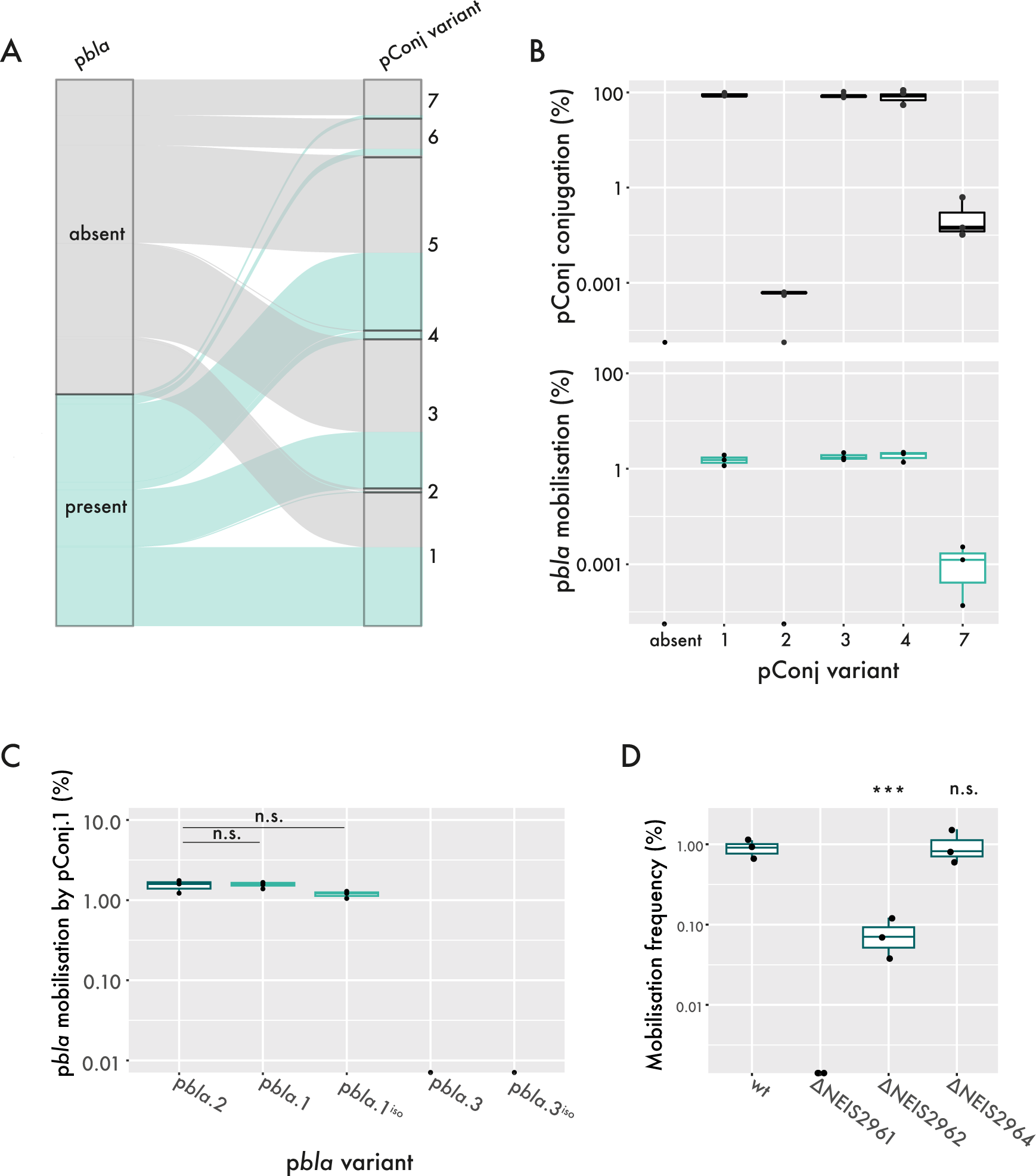
p*bla* mobilisation by pConj. (A) Sankey plot of pConj carrying isolates (n=4,883 isolates)^1^, displaying the presence of p*bla* (left) and co-occurrence of p*bla* with individual pConj variants (right). (B) Conjugation rates of pConj variants (top) and the mobilisation rates of co-located p*bla*.1 (bottom). (C) Mobilisation rates of wild type and isogenic p*bla* variants (p*bla*^iso^) by pConj.1. (D) The impact of single *mob* gene knockouts in p*bla*.2 on p*bla* mobilisation frequencies. All assays consist of three individual repeats and were analysed by one-way ANOVA with Tukey multiple comparisons; n.s. p>0.05, *** p<0.001

### The restricted distribution of p*bla*.3 is associated with its immobility

There are conflicting data about p*bla*.3 mobility^26-28^. Therefore, we assessed the mobilisation of wild-type p*bla* variants by pConj.1. Results demonstrate that wild-type p*bla*.1 and p*bla*.2 are mobilised efficiently, while p*bla*.3 mobilisation was not detected (Figure 2C). To assess whether p*bla* variant deletions are responsible for these differences, we introduced variant-specific deletions into p*bla*.2, generating the isogenic plasmids p*bla*.1^iso^ and p*bla*.3^iso^. Mobilisation of the isogenic plasmids did not differ from wild-type plasmids (p=0.82, Figure 2C), indicating the variant-specific deletions are responsible for mobilisation differences. Of note, the immobility of p*bla*.3 is evident from its restricted distribution in three related lineages, whilst p*bla*.1 and p*bla*.2 are found across the gonococcal population (Supplementary Figure 2)^1^.

Next, we examined the genes responsible for the immobility of p*bla*.3, and generated p*bla*.2 mutants lacking genes absent in p*bla*.3. Deletion of *mobA* (encoding the relaxase^29^) abolished p*bla* transfer (Figure 2D), indicating that the pConj relaxase cannot recognise p*bla oriT*. Removal of NEIS2962 significantly reduced p*bla* mobilisation (p=0.01, Figure 2D). NEIS2962 is related to MobC from *E. coli* plasmid RSF1010, which unwinds DNA at the *oriT*^30^. RSF1010. NEIS2962 homodimers are structurally related to MobC and predicted to recognise the p*bla oriT* but not a scrambled *oriT* sequence (Supplementary Figure 3). Deletion of NEIS2964 did not impact p*bla* transfer.

### TEM-135 confers increased penicillin resistance

Although p*bla*.3 is immobile, it is prevalent in Ng_cgc^400^s 25 and 298 (Table 1, Supplementary Figure 2)^1^. This suggests p*bla*.3, which carries TEM-135, confers a benefit to the gonococcus that has led to the clonal expansion of isolates with this plasmid. Therefore, we measured the penicillin MICs conferred by p*bla* variants. Whilst TEM-1 carrying p*bla*.1 and 2 conferred MICs of 8 µg/ml, p*bla*.3 with TEM-135 conferred a significantly higher MIC (32 µg/ml, p=0.003, Figure 3A). To establish whether the TEM variant determines resistance levels, we changed p*bla*.3 TEM-135 into TEM-1 by introducing a T182M substitution. This substitution reduced the p*bla*.3 MIC to levels of p*bla*.1/p*bla*.2, demonstrating that TEM-135 confers elevated MICs (Figure 3B). We also compared resistance conferred by TEM-1, TEM-1_P14S_ and TEM-135 (which together account for >95% of gonococcal TEMs^1^) expressed by p*bla*.2. Again, TEM-135 significantly increased MICs (128 µg/ml *vs.* 8 µg/ml with TEM-1, p<0.001, Figure 3C).

**Figure 3:**
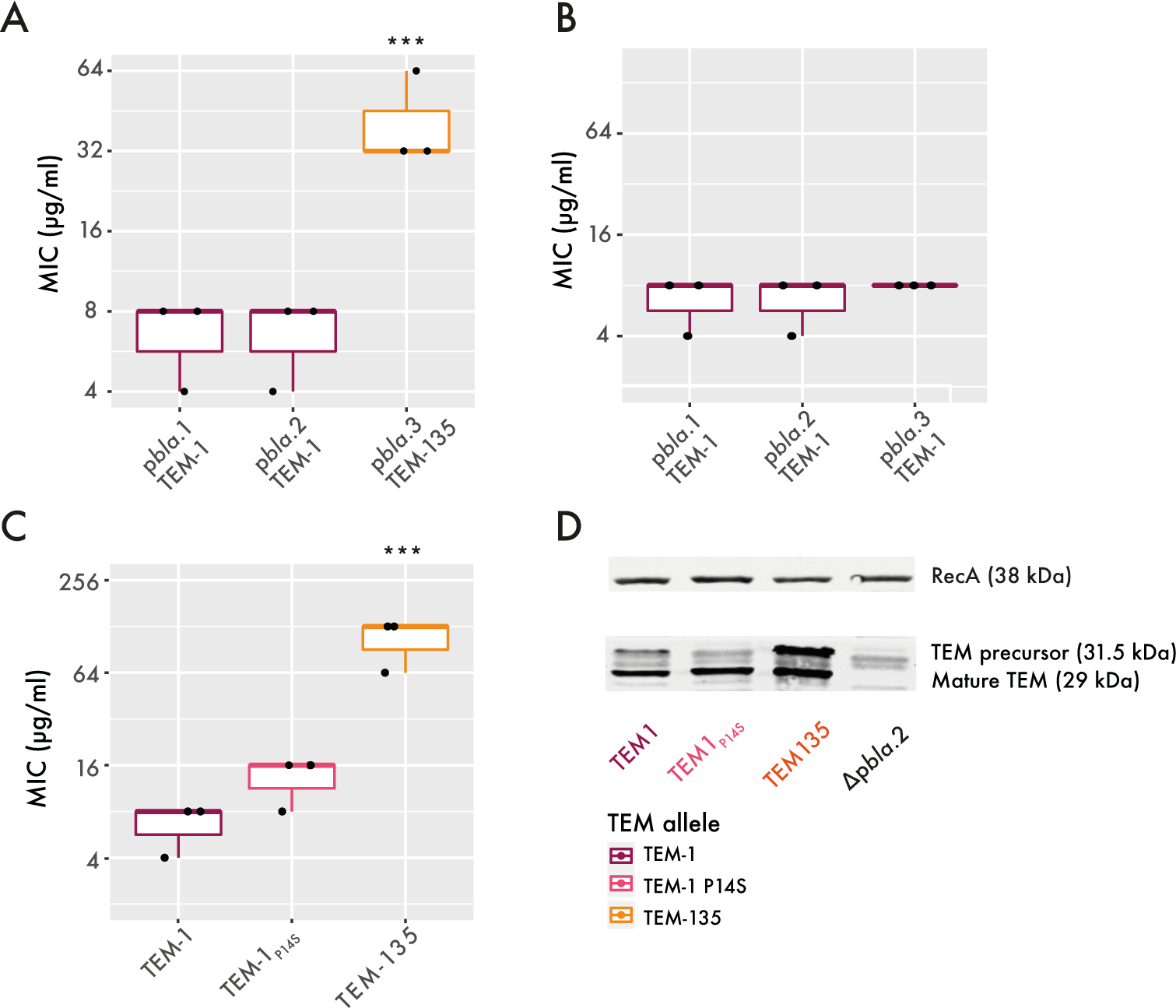
TEM-135 increases the MIC. (A) Penicillin G MICs of p*bla* variants in FA1090 isogenic strain background (one-way ANOVA on log_2_-transformed MIC values with Tukey multiple comparisons of means; *** p<0.001). (B) MICs of TEM-1 in different p*bla* variant backbones. (C) MICs of different TEM variants in p*bla*.2 backbone (one-way ANOVA on log_2_-transformed MIC values; *** p<0.001). (D) Representative Western blot of cellular enzyme levels of different TEM variants.

**Table 1:**
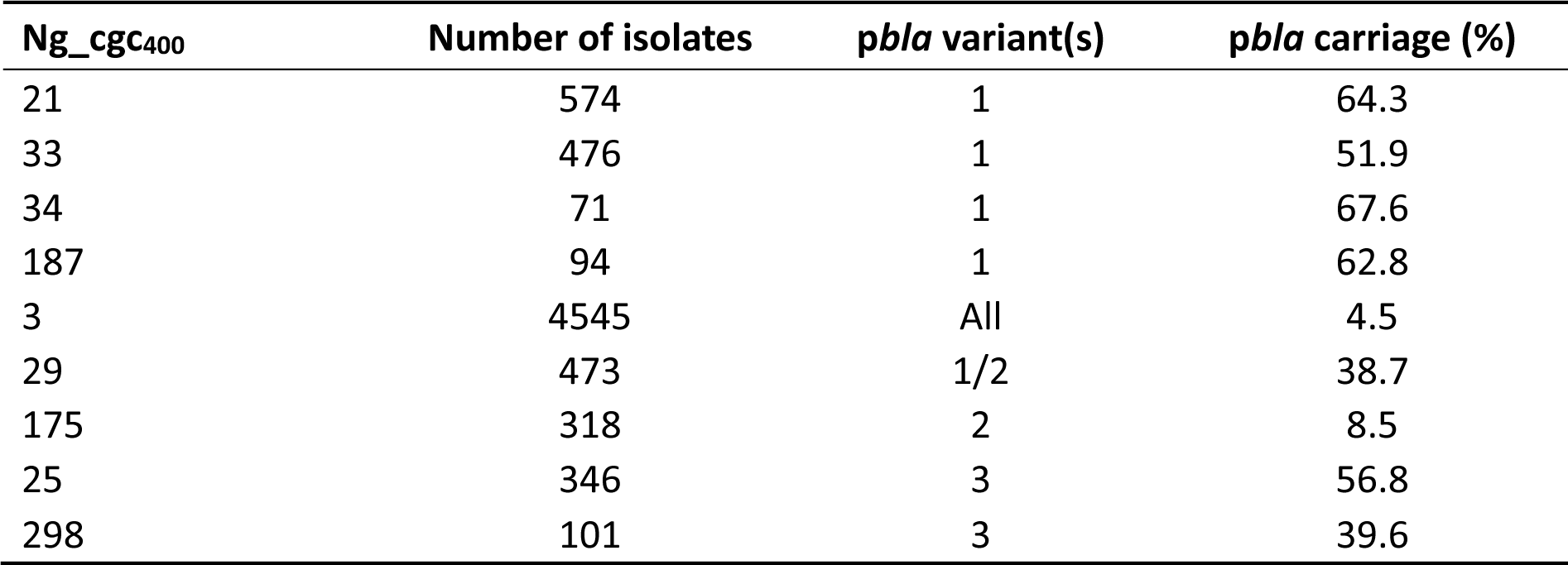
pbla prevalence in different Ng_cgc400. pbla-carrying lineages with >50 isolates and >5 isolates with pbla were assessed for their % pbla carriage.

To understand the basis for the different MICs, we assessed cellular TEM levels. Levels of mature TEM-135 (29 kDa) were significantly higher than TEM-1 or TEM-1_P14S_ (Figure 3D, Supplementary Figure 4), consistent with increased stability of TEM-135^17^. In conclusion, the appearance of TEM-135, particularly associated with p*bla*.3, provides a significant benefit to the gonococcus by enhancing resistance against beta-lactams, with MICs correlating with cellular TEM levels.

### p*bla* has evolved with reduced fitness costs

Plasmids often impose fitness costs, disadvantaging isolates which carry plasmids^31^. We therefore assessed the fitness costs of p*bla* by introducing p*bla*.1 into isolates from a range of lineages, and competing plasmid-carrying *vs.* plasmid-free strains. p*bla*.1 had no detectable fitness cost in any isolate (Figure 4A), consistent with its continued prevalence in the gonococcus (Figure 4B).

**Figure 4:**
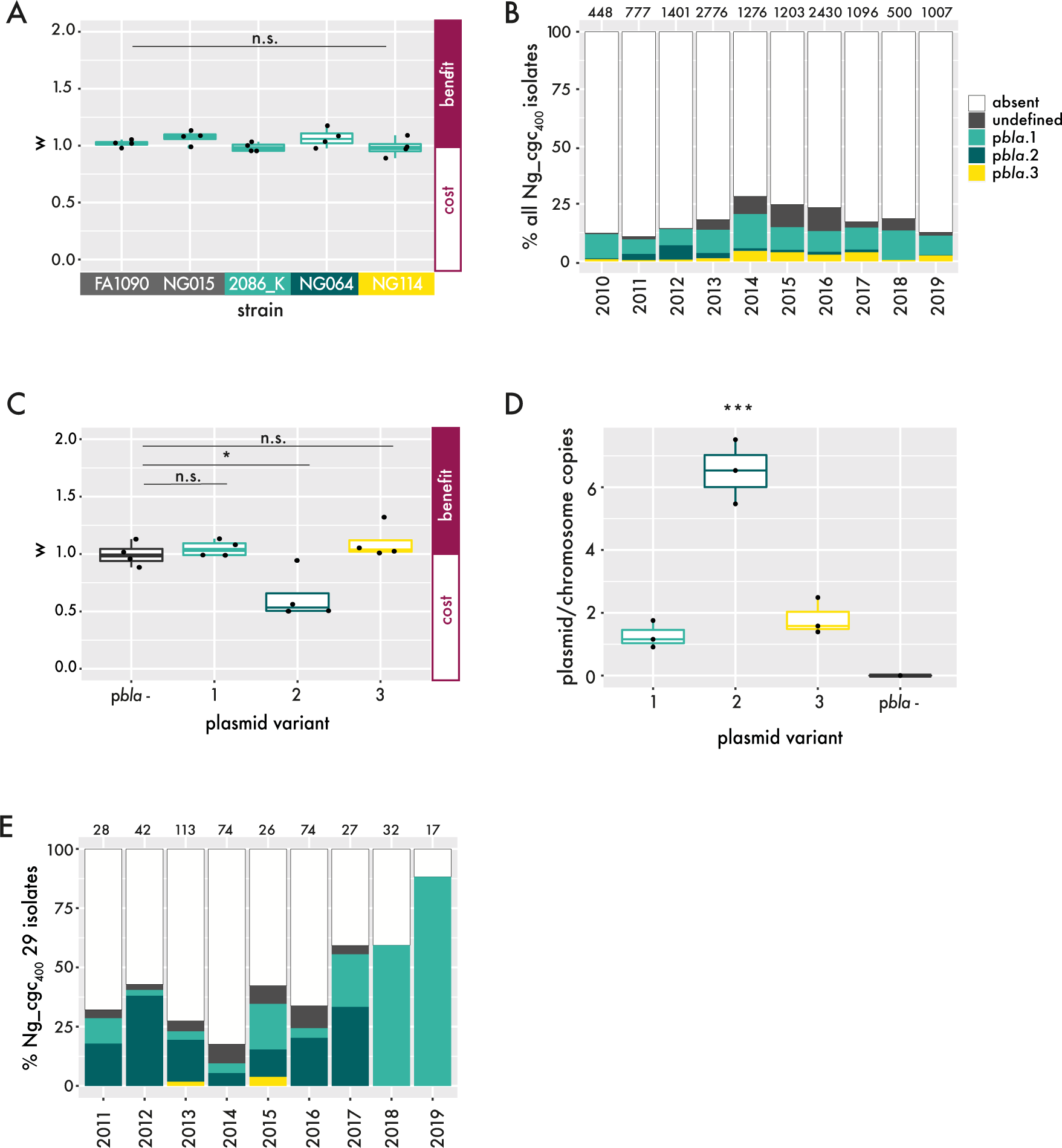
Impact of p*bla*-imposed fitness cost on its prevalence in the population. (A) Fitness cost (w) of p*bla*.1 in clinical isolates from different p*bla*-free (grey) or p*bla*-associated lineages (turquoise, p*bla*.1-associated; dark green, p*bla*.1/p*bla*.2-associated; yellow, p*bla*.3-associated). w>1 indicates a benefit, whereas w<1 signifies a cost of plasmid carriage. (B) Proportional p*bla* carriage in gonococcal isolates deposited on PubMLST between 2010 and 2019 (n=12,914 isolates). Colours show p*bla* variant carried and numbers above bars indicate the number of samples in the respective year. (C) Fitness cost of p*bla* variants in FA1090 isogenic strain background were assessed in four independent replicates (one-way ANOVA with Tukey multiple comparisons, n.s. p>0.05; * p<0.05). (D) Copy number of p*bla* in FA1090 isogenic strain background was assessed by ddPCR (one-way ANOVA with Tukey multiple comparisons; *** p<0.001). (E) p*bla* carriage in isolates from the p*bla*.1/p*bla*.2-associated Ng_cgc_400_ 29 between 2011 and 2019 (n=433 isolates). Bar colours indicate p*bla* variant and numbers above the bars specify the number of samples in the respective year.

In contrast to p*bla*.1 and p*bla*.3, p*bla*.2 inflicts a significant fitness cost (Figure 4C), and has a higher copy number (>6 *vs*. 1-2 plasmids/chromosome, Figure 4D). This could explain the decreasing prevalence of p*bla*.2 over time seen across all available WGS (Figure 4B). We also examined the prevalence of p*bla*.1 and p*bla*.2 within a single lineage (Ng_cgc_400_ 29 which harbours both variants) to account for potential sampling bias. Between 2010 and 2020, there has been a shift from p*bla*.2 to p*bla*.1 in this lineage (Figure 4E). Further evidence of relative success of the p*bla* variants is revealed by their abundance within a lineage. p*bla*.1 is prevalent in lineages, whilst p*bla*.2 is only present at low frequency in lineages (Table 1), reflecting its fitness costs and a lack of expansion of plasmid-carrying isolates within a lineage.

Taken together, fitness costs imposed by p*bla*.2 are consistent with its low prevalence in the gonococcal population compared with p*bla*.1. p*bla*.3 with TEM-135, which evolved from p*bla*.2, confers elevated penicillin resistance without fitness costs, and is associated with the success of a small group of related lineages.

## DISCUSSION

Plasmids are important vehicles for AMR, with resistance plasmids amongst the most diverse and mobile^32^. Here, we investigated the origin of the beta-lactamase plasmid p*bla* which is largely restricted to the gonococcus, a WHO priority pathogen. Our analysis indicates that p*bla* was acquired by *N. gonorrhoeae* at least twice from *Haemophilus*. Interestingly, *H. ducreyi* strains are divided into two clades, which diverged ∼1.9 million years ago^33^; p*bla.*1- and p*bla.*2-like plasmids are in Class I and Class II isolates, respectively. Thus, the independent acquisition of p*bla*.1 and p*bla*.2 by the gonococcus likely reflects separate events involving the two clades. Both intergenic transfers were accompanied by truncation of Tn*2*, suggesting the transposase is disadvantageous in the gonococcus.

In women, *N. gonorrhoeae* primarily causes cervicitis, while *H. ducreyi* causes ulcers at the vaginal entrance and cervix. In men*, N. gonorrhoeae* primarily causes urethritis and *H. ducreyi* mainly causes penile ulcers^34^. However, ∼3.5% of men with chancroid also have urethritis^34^. Therefore, these species can occupy the same niche, providing opportunities for gene transfer. Interestingly, p*bla* carriage is negligible in *H. influenzae* and *N. meningitidis* which inhabit the nasopharynx. p*bla* was not detected in any non-invasive *Neisseria* spp. (41,158 isolates, 30 species), and in only three of 39,372 *N. meningitidis* isolates in distinct lineages (ST-5, ST-11 and ST-32 complexes). This could reflect the renal excretion of beta-lactams^35^, favouring p*bla* carriage for bacteria inhabiting the urogenital tract compared with other sites. Of note, the meningococcal urethritis clade (NmUC) evolved from ST-11 *N. meningitidis* by acquiring genetic elements (but not p*bla*) from *N. gonorrhoeae* ^36,37^. We found p*bla* in an ST-11 meningococcal isolate, suggesting p*bla* could appear in NmUC.

Plasmids can be successful in bacterial populations by spreading into diverse lineages and/or through clonal expansion of plasmid-carrying isolates. We found that p*bla* is associated with pConj variants that can effectively spread p*<u>bla</u>* through the gonococcal population. Given the higher rates of pConj conjugation (>75%) compared with p*bla* mobilisation (∼1%), the spread of p*bla* into a lineage is likely to be accompanied by pConj, maintaining the close association between these plasmids.

The most frequent and widespread p*bla* variant, p*bla*.1, does not impose fitness costs and is mobilised efficiently by common pConj variants such as pConj.1. p*bla*.2 is also mobile but imposes fitness costs. Compared with p*bla*.1, p*bla*.2 has a second replication initiation protein and additional origins of replication (*ori2* and *ori3*)^1,38^. This might explain differences in copy number and fitness costs of p*bla*.1 and p*bla*.2; plasmid Rep proteins can sequester host DNA replication machinery^31^, causing fitness costs. We found that p*bla*.2 is present in lower prevalence in lineages than p*bla*.1, with evidence of a shift from p*bla*.2 to p*bla*.1 in a single lineage (Ng_cgc_400_ 29) over time. These observations are consistent with strains carrying p*bla*.2 being outcompeted by plasmid-free isolates or those with other p*bla* variants.

p*bla*.3-associated lineages have undergone clonal expansion indicating its successful adaptation to the gonococcus. Phylogenetic analysis indicates that TEM-135 originally arose in p*bla*.2. However, despite increased resistance levels conferred by TEM-135, the fitness cost of p*bla*.2 has likely undermined the success of TEM-135 in this p*bla* variant. We found that p*bla*.3 evolved from TEM-135 carrying p*bla*.2 through gene loss. This prevented the plasmid from being mobile, but with the trade-off of avoiding fitness costs. Consequently, p*bla*.3 has not spread in the gonococcal population but promoted the expansion of group of related lineages, likely though the increased beta-lactam resistance associated with TEM-135.

In summary, since emergence of gonococcal p*bla* in the 1970s, the evolutionary trajectory of this plasmid has been marked by its association with pConj variants that enable its spread through the population, the appearance of plasmid variants with minimal costs, and emergence of TEMs promoting higher resistance (*e.g.* TEM-135). A major concern is that the ESBL-permissive M182T substitution in TEM-135 is already widespread in gonococci^1,8^, especially in p*bla*.3. The intimate relationship between p*bla* and pConj highlights the threat posed by increased use of tetracyclines, as already witnessed in LMICs where gonococci have remarkably high plasmid carriage^7,39^. Similarly, the widespread implementation of Doxy-PEP will likely increase pConj and consequently p*bla* carriage, and promote the appearance of ESBL-expressing p*bla* with minimal fitness costs. This threatens treatment of cases and their contacts with third-generation cephalosporins, undermining the use of current first-line antibiotics used to control this important human pathogen.

## METHODS

### Analysis of *Haemophilus* spp. and *Neisseria* spp. genomes

Tn*2*-carrying isolates of *Haemophilus* spp. and *Neisseria* spp. were identified querying Tn*2* (GenBank accession: LC091537.1) against *Haemophilus* (accessed 24/07/2024, 4,640 isolates, 12 species) and *Neisseria* (accessed 24/07/2024, 41,158 isolates, 33 species) sequences in PubMLST^40^ (blastN word size: 11, scoring: reward: 2; penalty: -3; gap open: 5; gap extend: 2, sequence identity >99%, alignment length >50% query). p*bla*-like plasmids were confirmed by the presence of NEIS2960 (sequence identity >80%; alignment length >50%) and NEIS2358, and NEIS2961^1^.

### Plasmid variants

p*bla* and pConj were analysed in 15,532 gonococcal WGS (accessed 28/07/2022)^1,40^ with isolates from 1928-2022 and 66 countries. NEIS2220 indicated the presence of pConj, and its variants defined as previously^8^. Where the Ng_cp_5_ could not be assigned, plasmids were clustered with GrapeTree^41^, and variants assigned manually. p*bla* variants were typed using the Ng_p*bla*ST scheme^1^. For the population structure, isolates were grouped into core genome clusters according to the allelic profile of 1,668 core genes^42^; isolates were grouped with a cut-off of 400 allelic differences (Ng_cgc_400_).

### Phylogenetic analyses

A subset of 414 p*bla-*carrying isolates conserving the ratio of p*bla* variants (70% p*bla*.1, 14% p*bla*.2, 16% p*bla*.3, Supplementary Table 2)^1^ was selected to investigate the phylogenetic relationship of p*bla* variants. This included all p*bla*-containing isolates pre-dating 2000 (n=35). Isolates between 2000 and 2022 (n=379) were randomly selected using the r sample function^1^. Snippy v4.6.0 mapped plasmid reads to *H. ducreyi* DMC64 p*bla* (minimum coverage, 4 and base quality, 25). Multiple sequence alignments were generated with snippy-core v4.6.0/snippy-clean v4.6.0. Maximum likelihood trees were generated using RaxML-ng v1.2.2^43^ with 100 bootstrap replicates, rooted at *H. ducreyi* DMC64 p*bla*, and visualised with ape^44^ and ggtree^45,46^.

### Structure predictions

Analysis of NEIS2962 and RSF1010 MobC (GenBank accession: S96966.1) homodimers and NEIS2962 with p*bla oriT^29^* were performed using AlphaFold 3^47^ and PyMol v2.5.4^48^. Charge distributions were visualised with the Adaptive Poisson-Boltzmann Solver (APBS) electrostatics tool^49^.

### Bacterial strains/growth

Strains and plasmids used in this study are listed in Supplementary Tables 5 and 6, respectively. *E. coli* DH5α was grown on Luria-Bertani (LB) agar or in liquid LB shaking at 180 rpm. *N. gonorrhoeae* was grown on Gonococcal Base Media (GCB) agar plates or liquid media (GCBL)^50^ supplemented with 1% Vitox (Oxoid) at 37°C in 5% CO_2_. *H. ducreyi* was grown on chocolate agar plates supplemented with 1% IsoVitaleX at 35°C in 5% CO_2_. Antibiotics were added as follows: for *E. coli*, carbenicillin 100 μg/ml; for *N. gonorrhoeae*, carbenicillin 2.5 μg/ml; erythromycin 1 μg/ml; kanamycin 50 μg/ml, and tetracycline 2 μg/ml.

### Characterisation of *H. ducreyi* plasmids

Genomic DNA was isolated from *H. ducreyi* by harvesting bacteria from plates and the DNeasy Blood/Tissue Kit (Qiagen) with the modifications that cells were incubated in lysis buffer with 20 mg/ml lysozyme at 37°C for 2 hours and then proteinase K overnight at 56°C. Plasmids were analysed by Sanger sequencing.

### Transformation of gonococci

For electroporation, bacteria grown on GCB agar were resuspended in PBS (Sigma), adjusted to 5x10^7^ CFU/ml then washed three times with 20% glycerol / 1% MOPS (Sigma); electroporation was performed with 2.5 kV, 200 Ω, 25 mF. Cells were recovered in 1 ml of GCBL with 2% Vitox and plated on GCB agar. Plates were incubated for 3 hours, cells collected, and then transferred to selective media.

Δ*pilD*::*ermC* and Δ*pilD*::*aph* constructs were transformed into *N. gonorrhoeae* as described previously^23,25^. In brief, 1 μg of DNA was spotted onto plates, allowed to dry, and bacteria streaked over the spots. Plates were incubated for 8 hours, then bacteria were transferred onto selective agar. Transformants confirmed by PCR/Sanger sequencing.

### Plasmid modification

To generate p*bla*.1^iso^ and p*bla*.3^iso^, p*bla*.2 was cut with *Hin*dIII-HF and *Pvu*II-HF (NEB), respectively. p*bla*.1^iso^ was amplified with primers TE18/19 and PrimeSTAR GXL polymerase (Takara Bio); Gibson assembly was performed with primers TE20/21. p*bla*.3^iso^ was amplified in two fragments with TE7/TE17 and TE9/TE16. Plasmids were assembled using Gibson Hifi (NEB) and transformed into *E. coli* DH5*α*.

Point mutations in *bla*TEM were introduced using the RAIR method^51^. PCRs with primers (TE56/TE57, p*bla*.2 TEM-135; TE63/TE64, p*bla*.2^TEM-1 P14S^; TE65/TE66, p*bla*.3^TEM-1^) were performed using Herculase II polymerase (Agilent), with p*bla*.2 or p*bla*.3 as template. Products were purified (Promega Wizard PCR Clean-up) and transformed into *E. coli* DH5*α*.

*tetM*^+^ pConj.7 was constructed by amplifying *tetM* from *N. gonorrhoeae* WHO N using primers TE34/TE35. Flanking regions were amplified with primers TE36/37 and TE38/39, then joined by Gibson assembly; the product was amplified with TE36/39, then introduced into *N. gonorrhoeae* NG028 by transformation. All constructs were confirmed by sequencing.

### Conjugation and mobilisation assays

Donor (Δ*pilD*::*ermC*) and recipient (Δ*pilD*::*aph*) strains grown overnight were inoculated in 5 ml GCBL/1% Vitox at an OD_600_ of 0.1 and grown to mid-exponential phase (OD_600_ 0.6 - 0.8). The bacterial density was adjusted to 10^8^ CFU/ml and donor and recipient strains mixed in a 10:1 ratio. Bacteria (5 µl) were spotted onto GCB agar and incubated for 6 hours at 37°C, 5% CO_2_, harvested in 200 μl GCBL, then plated to GCB agar with antibiotics. Conjugation and mobilisation frequencies were defined as the number of transconjugants/recipients (n=3).

### Competition assays

Plasmids were introduced into FA1090 Δ*pilD*::*ermC* and competed against FA1090 Δ*pilD*::*aph* (n=4). Bacteria in PBS were adjusted to an OD_600_ 1, mixed 1:1, diluted to 10^5^ CFU/ml and added to 200 μl Fastidious Broth^52^ then grown at 37°C, 5% CO_2_, shaking at 180 rpm. After 24 hours, strains were enumerated by spotting on selective media. Fitness costs (w) were calculated by:

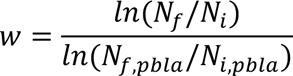

(w, relative fitness of p*bla*^+^ *vs*. p*bla^-^* strains; *N*_*i*_ and *N*_*f*_, p*bla^-^* strain at the beginning/end, respectively; *N*_*i*,*pbla*_and *N*_*f*,*pbla*_, same for p*bla*^+^ strain).

### Antibiotic susceptibility testing

Penicillin G MICs were assessed using the broth microdilution method^53^ in 96-well plates with 2-fold Penicillin dilutions in water (50 μl); strains grown overnight on GCB agar were resuspended in PBS (Sigma), then diluted in 2x FB/2% Vitox to 10^5^ CFU/ml. Bacteria (50 μl) were transferred into each well and incubated for 24 hours.

### SDS page and Western blot analysis

Bacteria were grown to mid-exponential phase, added to an equal volume of 2x SDS-PAGE buffer, run on 12% SDS-polyacrylamide gels, and transferred to Protan nitrocellulose membranes (GE Healthcare) using the Trans-Blot Turbo System (Bio-Rad). Membranes were blocked in PBS/0.5% Tween-20/5% milk, washed thrice and incubated with the primary antibodies (Rabbit anti-RecA, Abcam, ab63797, 1:5,000; Mouse anti-TEM, Abcam, 8A5.A10, 1:1,000) for 2 hours. After washing, membranes were incubated with secondary antibodies (LI-COR Biosciences, 925-68071 IRDye® 680RD Goat anti-Rabbit IgG and 925-32210 IRDye® 800CW Goat anti-Mouse IgG) at a final dilution of 1:10,000 for 1 hour, washed, then imaged using LI-COR Biosciences.

### Plasmid copy number

Copy number of *recA* and plasmid *tnpR* were quantified using the QX200 Droplet Digital PCR system (Bio-Rad) as described previously^54^. ddPCR contained 1x EvaGreen super mix (Bio-Rad), and TE79/TE80 (*recA*) or TE81/TE82 (*tnpR*). After thermal cycling, data were analysed using the QX200 Droplet Reader with QuantaSoft software (Bio-Rad).

### Statistics and data analysis

Data analysis was performed in R version 4.1.1 using base R and the tidyverse package^55^. Plots were generated with ggplot2^56^. A p value <0.05 was considered statistically significant.

## SUPPLEMENTARY FIGURES

**Supplementary Figure 1:**
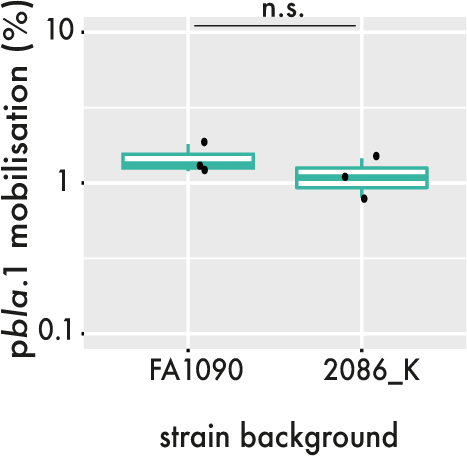
p*bla.*1 mobilisation in isogenic matings with the *N. gonorrhoeae* strains FA1090 and 2086_K (Welch two-sample t-test, p=0.66).

**Supplementary Figure 2:**
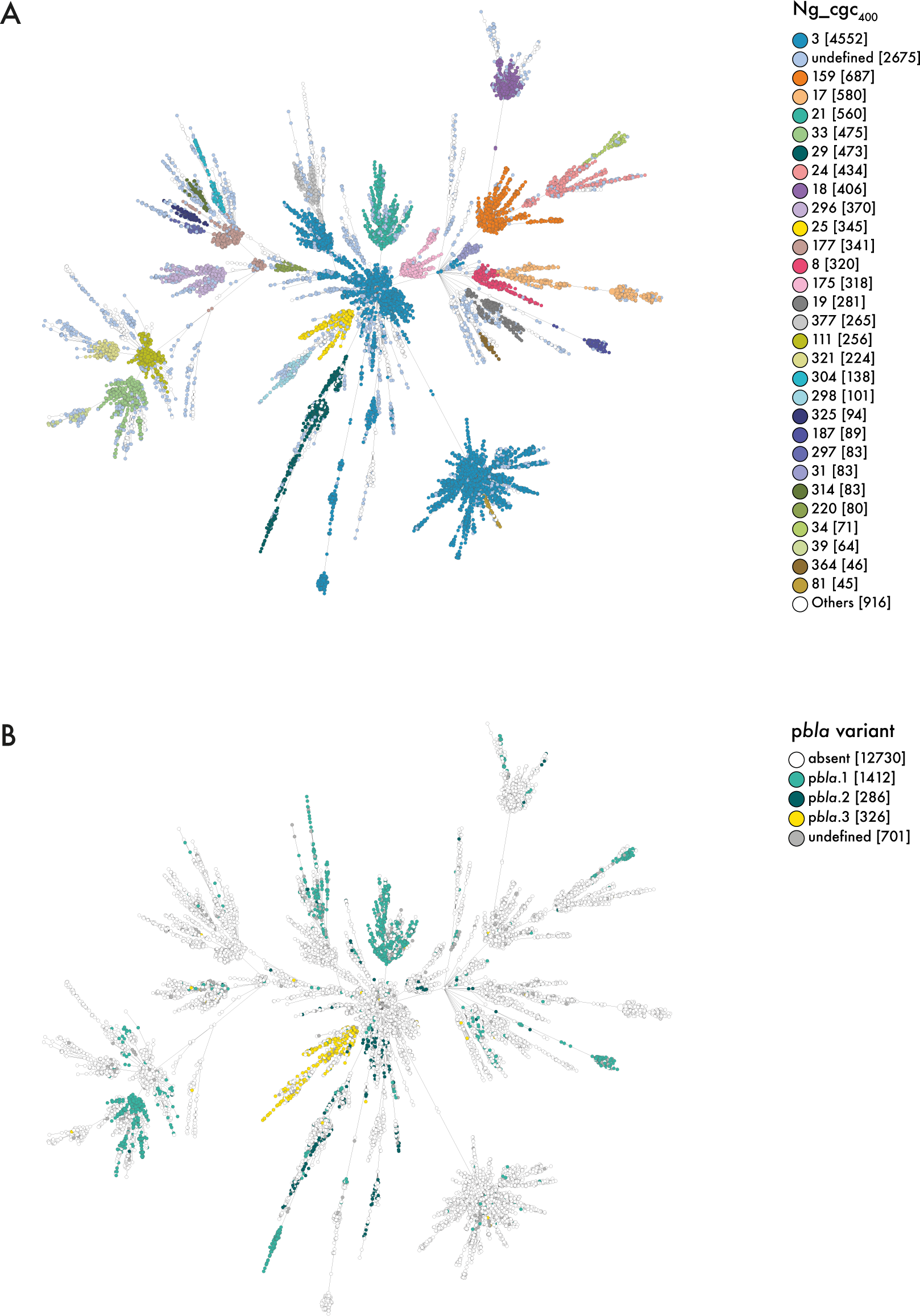
Minimum spanning trees of *N. gonorrhoeae* clustered by core genome allelic differences with distribution of p*bla* variants. Individual dots represent isolates that are coloured according to Ng_cgc_400_ (A) or p*bla* variant carried (B).

**Supplementary Figure 3:**
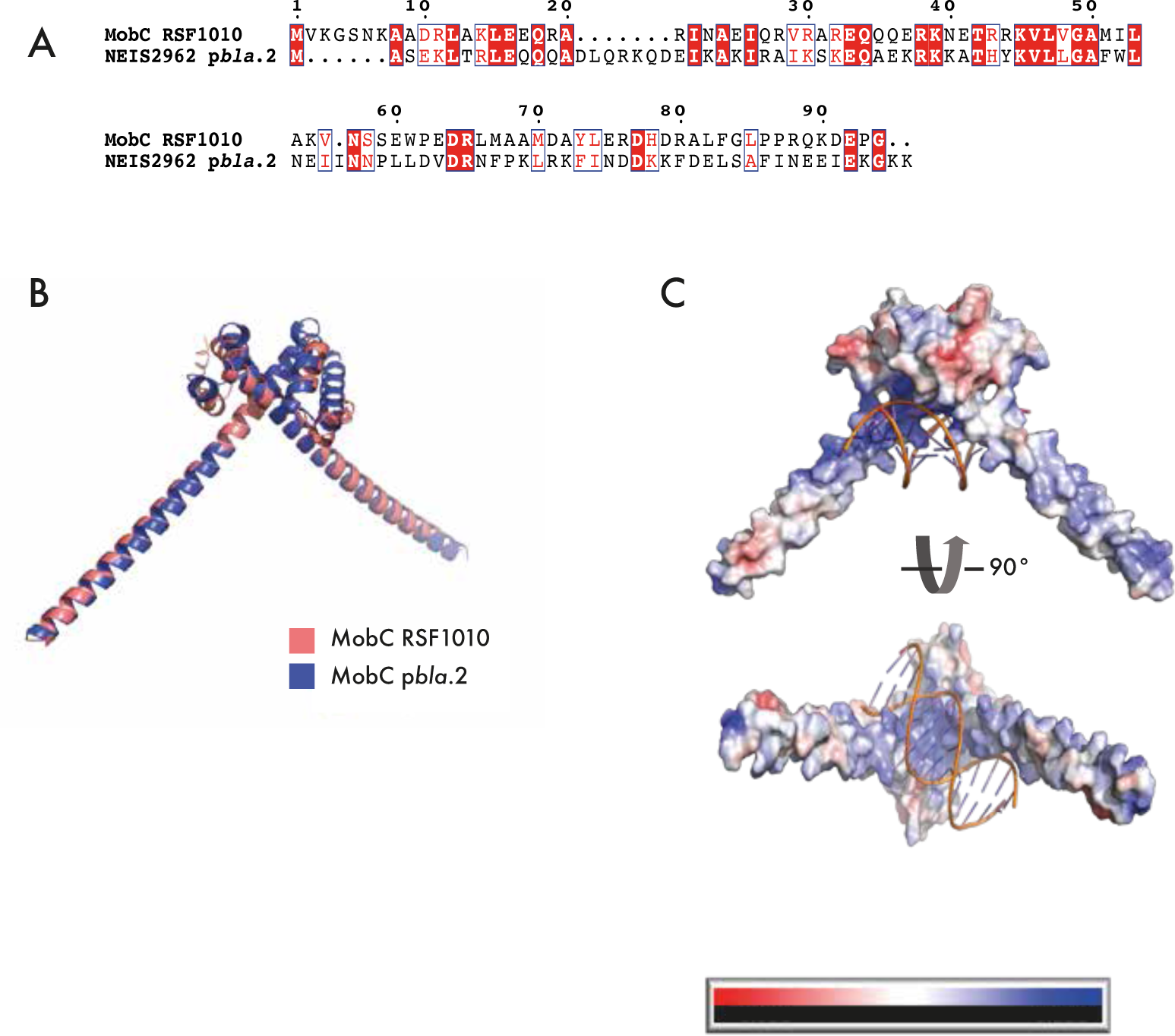
(A) Alignment of MobC from *E. coli* plasmid RSF1010 (Genbank accession: S96966.1) and NEIS2962 from p*bla*.2 (Genbank accession: NZ_LT591911). Amino acid sequences were aligned with COBALT^3^ and the alignment visualised with ESprit^9^. Identical residues are shown in white on red background, residues with a similarity score >0.7 are framed in blue and the remaining residues are shown in black. (B) Superimposed AlphaFold structure prediction of MobC from the *E. coli* plasmid RSF1010 (salmon, Genbank accession: S96966.1) and NEIS2962 (blue) dimers (Match Align: 677.7, RMSD: 0.775Å) (C) Electrostatics prediction of NEIS2962 homodimer with *oriT* sequence using the Adaptive Poisson Boltzman Solver Electrostatics Plugin. Negatively and positively charged regions are shown in red and blue, respectively.

**Supplementary Figure 4:**
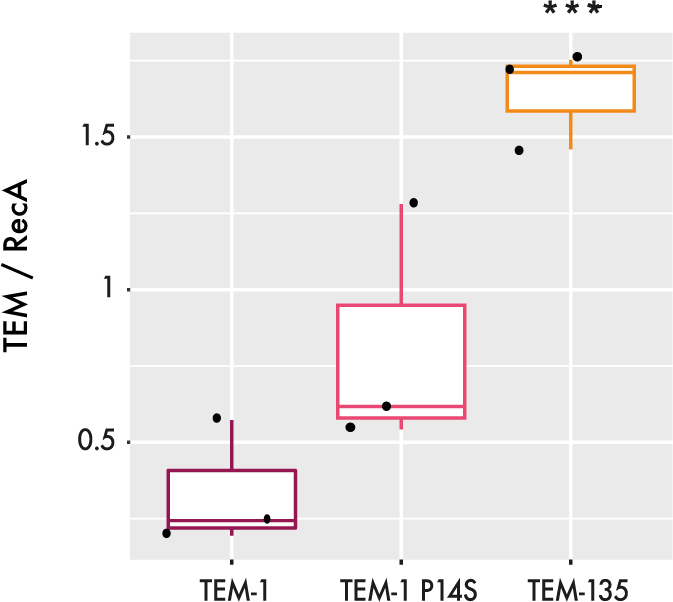
Cellular levels of different TEM variants in an isogenic FA1090 background. TEM/RecA ratios of whole cell lysates were visualised by Western blot analysis and quantified with the LI-COR system (one-way ANOVA with Tukey multiple comparisons, n.s. p>0.05; *** p<0.001).

